# TREVI^XMBD^: A Transcriptional Regulation-driven Variational Inference Model to Speculate Gene Expression Mechanism with Integration of Single-cell Multi-omics

**DOI:** 10.1101/2023.11.22.568363

**Authors:** Lan Cao, Wenhao Zhang, Feng Zeng, Ying Wang

**Author notes:** To whom correspondence should be addressed: Feng Zeng,;, Ying Wang.

## Abstract

Single-cell multi-omics technology enables the concurrent measurement of multiple molecular entities, making it critical for unraveling the inherent gene regulation mechanisms driving cell heterogeneity. However, existing multi-omics techniques have limitations in capturing the intricate regulatory interactions among these molecular components. In this study, we introduce TREVI^XMBD^ (Transcriptional REgulation-driven Variational Inference), a novel method that integrates the well-established gene regulation structure with scRNA-seq and scATAC-seq data through an advanced Bayesian framework. TREVI^XMBD^ models the generation of gene expression profiles in individual cells by considering the integrated influence of three fundamental biological factors: accessibility of cis-regulatory elements regions, transcription factor (TF) activities and regulatory weights. TF activities and regulatory weights are probabilistically represented as latent variables, which capture the inherent gene regulatory significance. Hence, in contrast to gene expression, TF activities and regulatory weights that depict the cell states from a more intrinsic perspective, can keep consistent across diverse datasets. TREVI^XMBD^ exhibits superior performance when compared to baseline methods in a variety of biological analyses, including cell typing, cell development tracking, and batch effect correction, as validated through comprehensive benchmarking. Moreover, TREVI^XMBD^ can reveal variations in TF-gene regulation relationships across cells. The pretrained TREVI^XMBD^ model can work even when only scRNA-seq is available. Overall, TREVI^XMBD^ introduces a pioneering biological-mechanism-driven framework for elucidating cell states at a gene regulatory level. The model’s structure is adaptable for the inclusion of additional biological factors, allowing for flexible and more comprehensive gene regulation analysis.

## 1. Introduction

Living organisms are intricately regulated by a multitude of genetic elements, including transcription factors (TFs), cis-regulatory elements (CREs), and the accessibility of chromatin regions. Deciphering this multifaceted gene regulation is of great importance. Multi-omics sequencing techniques have emerged as a promising approach to address this challenge. They enable the simultaneous profiling of cell transcriptomes and chromatin accessibility using single-cell RNA sequencing (scRNA-seq) and single-cell Assay for Transposase-Accessible Chromatin with sequencing (scATAC-seq), respectively.

The inference of gene regulation from single-cell multi-omics data typically involves the integration of distinct modality data, followed by the analysis of gene regulation networks. However, multi-omics data are susceptible to various sources of noise. To mitigate noise and improve the identification of cell states, several integration methods have been developed, encompassing shared information across distinct omics and advanced graph structures. Prominent examples include MultiVI [1], Cobolt [2], scMVP [3], and GLUE [4]. In the context of each cell state, the inference of TF-gene regulatory relationships is accomplished by assessing the possible binding of TFs to CRE regions of genes, as exemplified by DIRECT-NET [5], or by employing association analysis between expression profiles of TFs and genes, as demonstrated by scGRNom [6]. However, the division between cell state definition and regulatory inference can introduce biases at each step, limiting the accurate representation of the dynamic and diverse nature of regulatory mechanisms governing cell fate decisions. Furthermore, multi-omics techniques face challenges in directly capturing physical contacts among genetic elements, making it challenging to infer gene regulation from these multiomics data solely through ‘guilt by association’ approaches.

To address these challenges, we develop TREVI^XMBD^, an innovative variational inference model. It devises a Bayesian framework to incorporate the well-established gene regulation structure as prior knowledge and facilitate the integration of multi-omics data and the inference of gene regulation simultaneously. TREVI^XMBD^ aims to optimize the estimation of TF activities and the TF-gene interactions by precisely modeling the generation of single-cell profiles under the synergistic control of TFs and other genetic elements. We constructed seven benchmarking datasets, covering diverse cell compositions and temporal omics profiles, to validate the effectiveness of TREVI^XMBD^. Across various analyses, including cell typing, batch effect correction and cell development tracking, TREVI^XMBD^ consistently outperformed the baseline methods. Additionally, TREVI^XMBD^ enables the construction of a TF-gene regulatory network that reveals cell state-specific dynamics. Furthermore, TREVI^XMBD^ can be pre-trained with multi-omics data and applied to scRNA-seq alone, showcasing its broad applicability. Overall, TREVI ^XMBD^ provides novel insights driven by biological mechanisms for characterizing cell states. The structure of TREVI ^XMBD^ is adaptable for the inclusion of additional biological factors, allowing for flexible and more comprehensive gene regulation analysis.

## 2. Methods

### 2.1 Framework of TREVI^XMBD^

As illustrated in Figure 1A, the regulation of gene expression is a multifaceted process. Initially, chromatin regions become accessible, allowing for the binding of TFs to CREs. Subsequently, TFs engage with these regions, initiating gene expression. It is reasonable to postulate that the extent of gene expression can be predicted based on the characteristics of chromatin accessibility and TF activity. On one hand, regions of chromatin with increased accessibility are more likely to attract TF binding. On the other hand, TFs with higher activity levels exhibit a greater affinity for CREs, signifying enhanced regulatory control over the target gene. Drawing inspiration from these principles, TREVI^XMBD^ comprises three distinct components designed to model chromatin accessibility, TF activity, and TF-gene regulation, respectively (Fig. 1A). Detailed explanations of each component are presented below.

**Figure 1.**
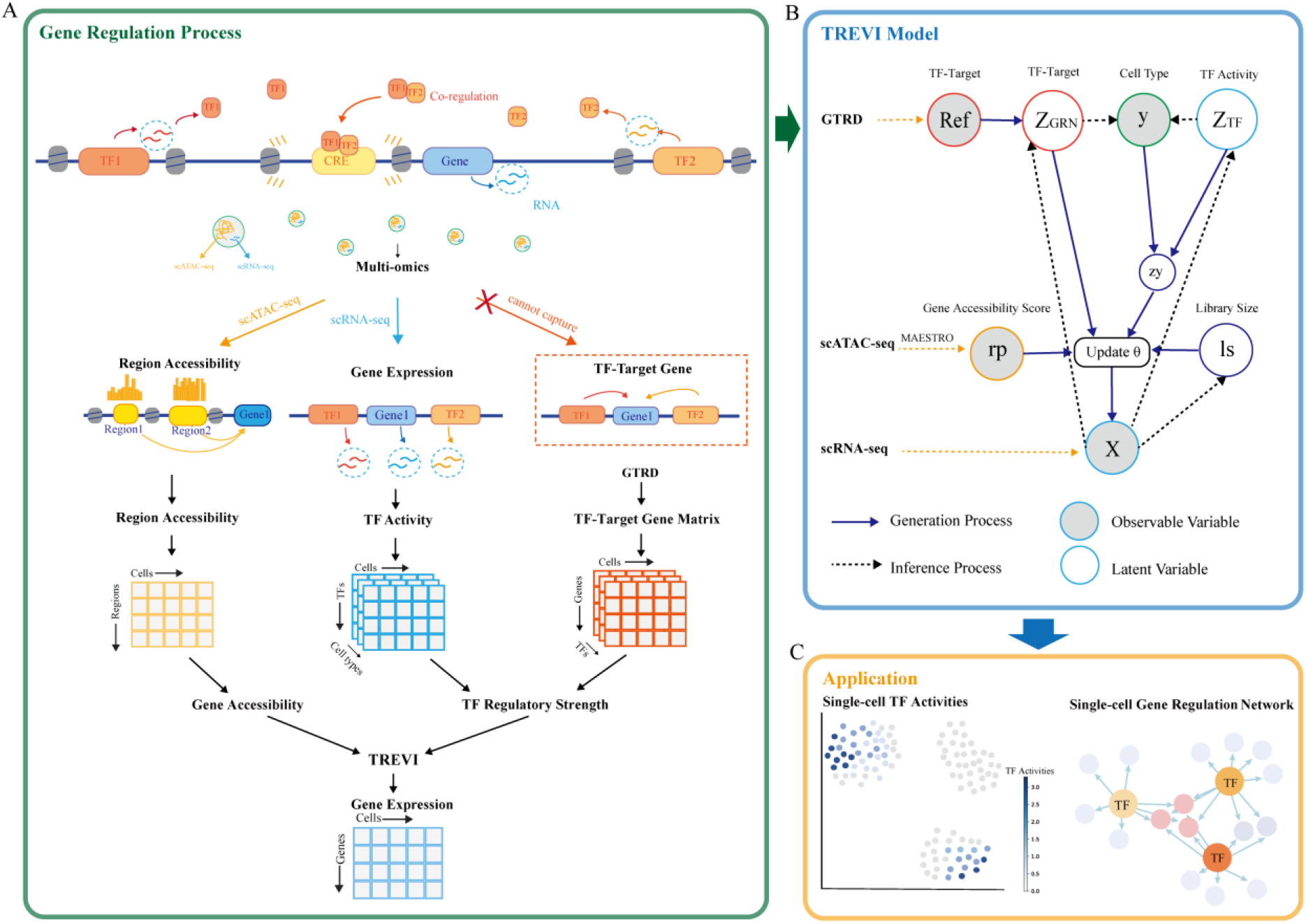
Framework of TREVI^XMBD^. TREVI^XMBD^ is built on the premise that gene expression is regulated by several key factors: TF activity, TF-gene regulating weight, and chromatin accessibility. It leverages multi-omics data to learn the latent variables, TF activity and TF-gene regulatory weight within cells. TREVI^XMBD^ triggers the generation process for gene expression profile and infers the latent variables on scRNA-seq and scATAC-seq data. The derived TF activity can be applied to identify cell states, explore cell-type-specific TFs, and correct batch effects. Similarly, the inferred TF-gene regulatory weights facilitate the construction of cell type-specific regulatory networks.

The first component focuses on the formulation of cell state-specific TF activity. TREVI^XMBD^ employs a latent variable model to describe TF activities. It collaborates with the TF-gene regulatory matrix, introduced subsequently, to generate expression profiles for target genes. Moreover, the TF activities derived from this latent variable model provide a more accurate characterization of cell states, extending beyond mere gene expression levels. The second component centers on calculating the accessibility of CRE regions. We define the probability of a gene being regulated by TFs by aggregating the accessibility scores of multiple CREs located within a 150-kilobase pairs proximity to the gene, following the protocol of MASTERO [7]. The third component aims to model the regulatory weights of each TF to its target genes. TREVI^XMBD^ employs a Bayesian approach to model these weights. It constructs the prior distribution by utilizing the regulatory interaction network between 2,986 TFs and their target genes derived from Gene Transcription Regulation Database (GTRD v19.04) [8].

These components are integrated into a Bayesian framework. The pre-processing of the raw data is described in Supplementary Materials S1. As shown in Figure 2B, TREVI^XMBD^ consists of a generation process and a variational inference process. Specifically, the generation process involves gene accessibility score, TF activity, and TF-gene regulatory matrix to generate gene expression profiles of individual cells based on zero-inflated multinomial distribution. The variational inference process is employed to infer the distribution parameters of the latent variables from the observable gene expression.

**Figure 2.**
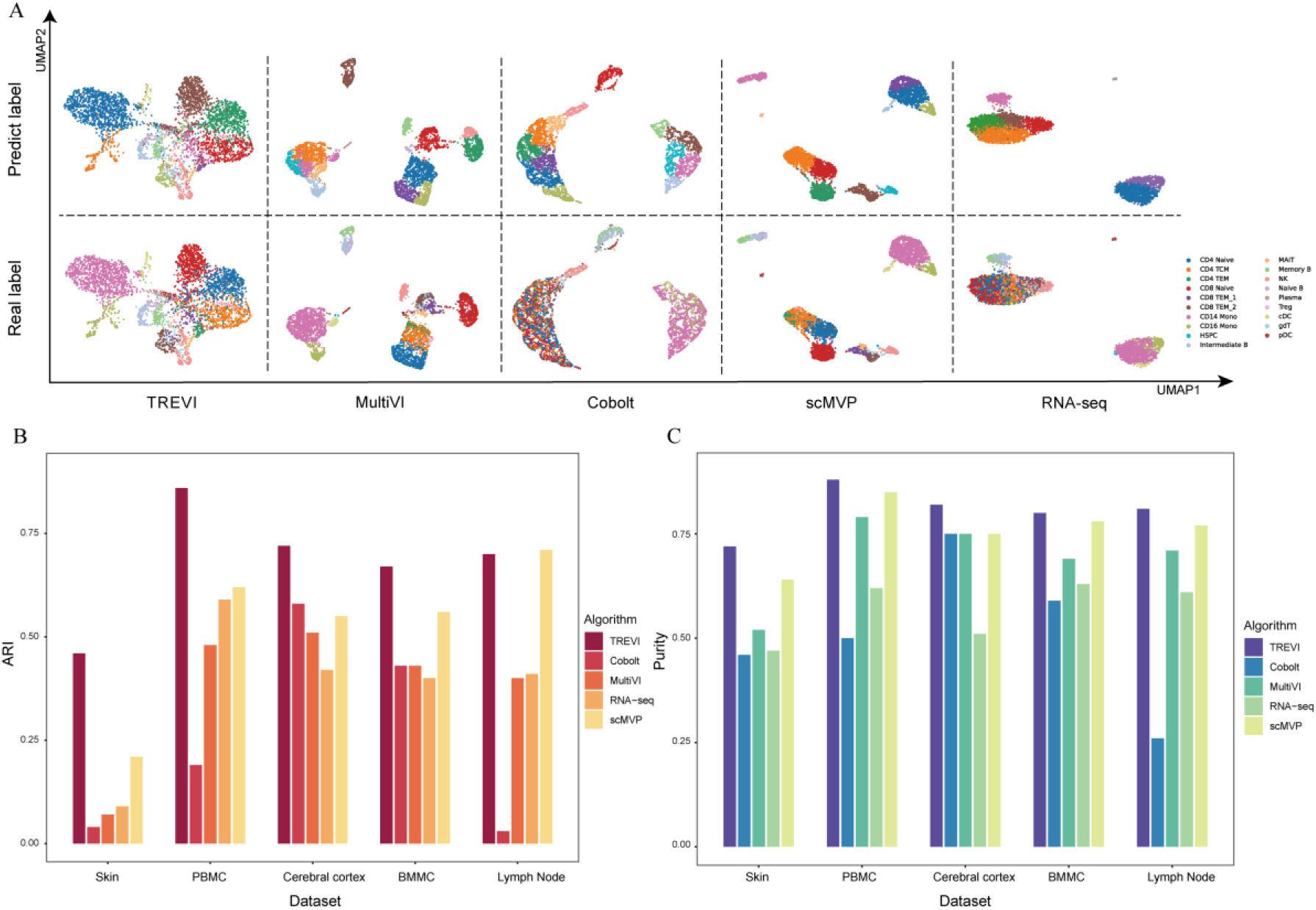
Performance comparison in cell type clustering. A. Visualization of clustering results between TREVI^XMBD^ and baseline methods. The two lines represent the predicted and true labels under five distinct methods applied to the PBMC dataset, respectively. It is observed that TREVI^XMBD^ effectively disperses the latent variables in accordance with the distributions of various cell types. B. The ARI values achieved by TREVI^XMBD^ and baseline methods across five datasets. C. Purity scores by TREVI^XMBD^ and baseline methods across five datasets.

### 2.2 Objective function

Let’s define symbols first. We use the variable *x* to represent the gene expression profile of individual cell. The inferred TF activity, the accessibility of the chromatin regions, and the TF-gene regulatory matrix are denoted as *z*_*TF*_, *rp* and *z*_*GRN*_ respectively. In addition, *x* is also controlled by *z*_*y*_ under the specific cell type *y* and the library size *ls* during the sequencing process.

For the data-generation process, the joint probability of all variables can be formulated as follows:

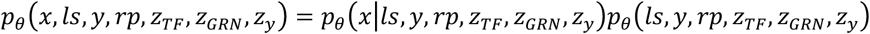

where the variable *y* denotes cell type and *ls* denotes library size. The variable *θ* represents the set of the parameters in this model. The likelihood function *p*_*θ*_(*x*| *ls, y, rp, z*_*TF*_, *z*_*GRN*_, *z*_*y*_) describes the mapping from latent regulatory factors to gene expression profiles through a nonlinear function implemented by deep neural networks. The marginal likelihood for *x* can be computed as follows:

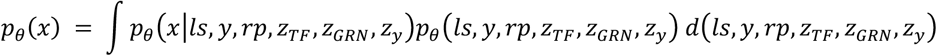

The parameters *θ* are estimated by solving the objective function argmax log(*p*_*θ*_(*x*)) . However, *p*_*θ*_(*x*) is intractable when latent variables are continuous. Therefore, we optimize the parameters in TREVI^XMBD^ through variational inference.

### 2.3 Generative process

The generation process of gene expression profile *x* is given here, beginning with the initialization of latent variables. TF activity variable *z*_*TF*_ initially follows a standard normal distribution:

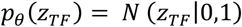

Cell type *y*, initially follows a One-Hot Categorical distribution:

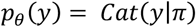

The latent variable *z*_*y*_ conforms to a normal distribution with parameters derived from *y* and *z*_*TF*_ though a neural network function *f*_*θ*_:

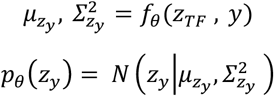

Gene accessibility score *rp* acts as the prior information extracted from the regulatory score matrix *R*.

The latent variable *z*_*GRN*_ that reflects the actual regulatory weights between TF and gene is also generated based on the prior knowledge *Ref*_*TF*−*gene*_ that deposits the known associations between TF and target genes, as follows:

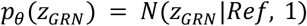

where regulatory relationships variable *z*_*GRN*_ initially follows a normal distribution with a mean of *Ref*_*TF*−*gene*_ .

Additionally, the library size *ls* is embraced, which follows a *Weibull* distribution [9]:

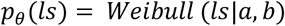

where the shape parameter *a* influences the skewness of the *Weibull* distribution, and *b* is the scale parameter that controls library size units.

We assume that gene expression profile *x* follows a *zero-inflated multinomial* distribution as follows:

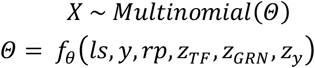

here *Θ* represents the parameter of the *zero-inflated multinomial* distribution, which is obtained through a neural network function *f*_*θ*_.

Taken together, the generation process of gene expression profile *x* can be formulated as follows:

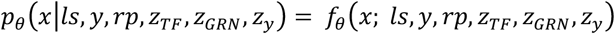

where *f*_*θ*_(*x*; *ls, y, rp, z*_*TF*_, *z*_*GRN*_, *z*_*y*_) represents the hierarchical nonlinear transformations.

### 2.4 Inference process

The generation process describes the probabilistic generation of gene expression profile *x* based on the regulatory variables. In the inference process, the real distribution parameters in the generation process are inferred based on the observable data. *q* represents the distributions we constructed with neural networks.

Specifically, we use the constructed distribution *q*_*φ*_(*z*_*TF*_|*X*) to infer TF activity *z*_*TF*_ from the observed gene expression profile *X* derived from scRNA-seq, as follows:

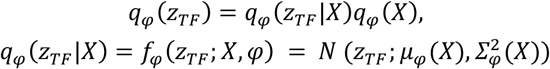

where *φ* is the set of parameters in the constructed distributions, *μ*_*φ*_(*X*) and 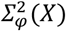 are directly obtained from the observed data *X, f*_*φ*_ (*z*_*TF*_; *X, φ*) is the nonlinear transformation that map *z*_*TF*_ from *X*.

Similarly, we use *q*_*φ*_(*z*_*GRN*_|*X*) to infer the TF-gene regulatory weights from gene expression profile *X*, and *q*_*φ*_(*ls*|*X*) to infer the library size *ls* from gene expression profile *X*, which are formulated as follows:

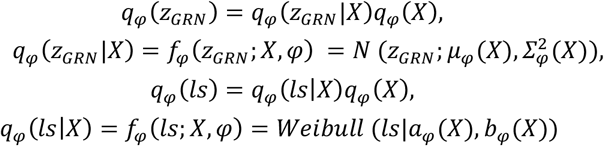

*q*_*φ*_(*y*|*z*_*TF*_ ) and *q*_*φ*_(*y*|*z*_*GRN*_) are used to infer the cell type labels *y* from *z*_*TF*_ and *z*_*GRN*_, as follows:

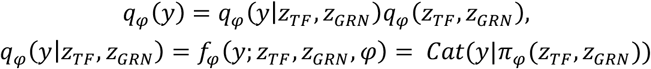

During the inference process, the used distributions can extract latent information and parameters from the observed data, providing prior support for the generation process.

The loss function and parameter optimization are described in Supplementary Materials S2.

## 3. Results

### 3.1 Datasets

To evaluate TREVI^XMBD^, we utilized seve<u>n</u> datasets. These datasets encompassed both multi-omics and single-omics data, comprising paired scRNA-seq and scATAC-seq data, as well as standalone scRNA-seq data. Summaries about these datasets are given in Supplementary Materials S3. These datasets represent different biological scenarios. Specifically, five of the datasets originated from various tissue compartments, including Peripheral Blood Mononuclear Cell (PBMC) [4], Bone Marrow Mononuclear Cell (BMMC) [10], Lymph node [11], Skin [12], and Cerebral cortex [13]. They were used to evaluate TREVI^XMBD^’s performance in cell typing and batch effect correction. Additionally, we employed a longitudinal time-series dataset, denoted as A549 [14], to evaluate TREVI^XMBD^’s capability to distinguish between different stages of cell evolution. The final dataset, PBMC-IFNB [15], was employed to evaluate the generalization ability of TREVI^XMBD^ as a pre-trained model. This dataset consisted of scRNA-seq data from both normal PBMCs and PBMCs subjected to IFNB stimulation. The cell type annotations are provided in dataset annotations, some of the annotations were obtained with the Seurat method [16] as indicated in the dataset. Detailed descriptions of the performance evaluation metrics can be found in Supplementary Materials S4.

### 3.2 TF activities improve the characterization and classification of cell types

First, we assessed the efficacy of TF activities for the characterization and classification of distinct cell types. We applied TREVI^XMBD^ to derive the low-dimensional representations based on TF activities from five datasets, including PBMC, BMMC, Lymph node, Skin, and Cerebral cortex. For comparative analysis, we also leveraged cell embeddings obtained through integrated multi-omics data using tools such as MultiVI, Cobolt, and scMVP, for more detailed introductions, please refer to Supplementary Materials S5. As a reference, we also created low-dimensional cell embeddings from scRNA-seq data alone using the standard UMAP algorithm within Seurat. Subsequently, we employed the Louvain algorithm [17] to identify cell clusters and compared clustering results and cell type annotations. Performance evaluation was conducted using established metrics, including Adjusted Rand Index (ARI) and Purity.

Our findings, as shown in Figure 2A, demonstrate that TREVI^XMBD^ effectively unveiled distinct cell clusters that aligned well with cell type annotations. Visual inspection of the low-dimensional embeddings in the PBMC dataset revealed highly separable cell clusters, consistent with the evaluation metrics presented in Figures 2B and 2C. In terms of ARI, TREVI^XMBD^ outperformed the baseline methods across four datasets, with a minor exception on the Lymph node dataset where it performed slightly worse than scMVP. Regarding Purity, TREVI^XMBD^ consistently exhibited superior performance across all five datasets when compared to baseline methods. This experiment underscores the effectiveness of TREVI^XMBD^ in inferring cell type-specific TF activities, which offer a more precise characterization of cell states in comparison to conventional gene expression profiles and integrated multi-omics data.

### 3.3 TF activity improves the characterization of temporal variability in perturbed cells

Next, we investigated the capacity of TREVI^XMBD^ to capture subtle temporal dynamics in perturbed cells of the same cell type, aiming to discern nuanced changes. For this purpose, we utilized the A549 dataset, comprising cells derived from the human non-small cell lung cancer (NSCLC) cell line, treated with Dexamethasone (DEX), and sampled at three distinct time points (0, 1, and 3 hour) [18]. Employing multi-omics sequencing, the dataset probed the impact of DEX on the evolution and reprogramming of human NSCLC cells, a process characterized by gradual, subtle alternations in cellular genetic elements. Our goal was to evaluate the capability of these methods to accurately capture time-induced nuanced dynamics while effectively mitigating the influence of unrelated signals, such as technical noise inherent in multi-omics data.

In this experimental setup, the TF activities inferred by TREVI^XMBD^ were employed to characterize cells at different time stages, in comparison to the baseline methods detailed earlier. As shown in Figure 3A, TREVI^XMBD^ effectively separated cells into distinct groups, each faithfully representing the respective time stage. Notably, these groups formed a chronological sequence in exact alignment with their collection times. In contrast, the baseline methods struggled to capture the temporal variations associated with the DEX perturbation. Specifically, their results exhibited a mingling of cells from different stages, rendering them indistinguishable.

**Figure 3.**
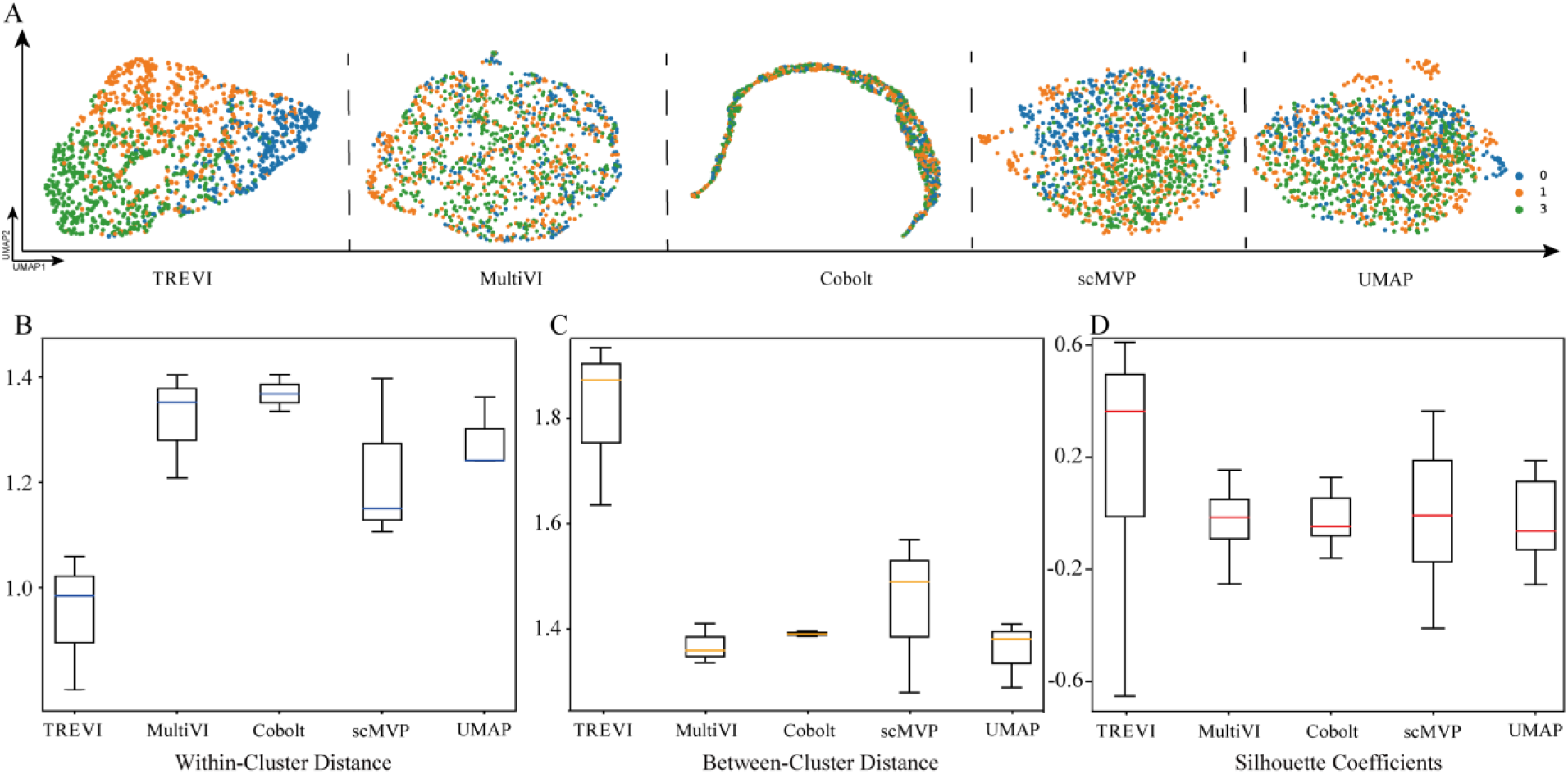
Performance comparison in capturing temporal variation of cells. A. The visualization of clustering results on A549 dataset between TREVI^XMBD^ and baseline methods. TREVI^XMBD^ effectively clusters cells on distinct time stages and follows correct time orders after drug perturbation. B. The boxplot of the within-cluster distance for five methods. C. Boxplot of the between-cluster distance for five methods. D. The boxplot of the Silhouette coefficient.

For a quantitative performance evaluation, we computed both within-cluster and between-cluster distances among cells collected at various time points. The within-cluster distance assessed the degree to which cells collected at the same time point cluster together, while the between-cluster distance measured the distinctiveness of cells collected at different time points. As shown in Figure 3B-C, TREVI^XMBD^ significantly reduced within-cluster distances compared to MultiVI (*p* − value = 2.87 × 10^−72^), Cobolt (*p* − value = 1.50 × 10^−124^), scMVP (*p* − value = 6.31 × 10^−38^) and UMAP (*p* − value = 8.29 × 10^−57^), indicating an effective grouping of cells collected at the same time point. Additionally, TREVI^XMBD^ significantly increased between-cluster distances compared to MultiVI (p − value = 1.04 × 10^−132^ ), Cobolt ( p − value = 4.38 × 10^−151^ ), scMVP ( p − value = 1.82 × 10^−88^ ), and UMAP ( p − value = 2.07 × 10^−134^), suggesting an effective separation of cells collected at different time points. These results were obtained through *Wilcoxon* rank-sum tests and corrected using Holm’s adjustment method [19].

Furthermore, we calculated the Silhouette coefficient to quantify the cohesion of cells within their respective clusters and their separation from other clusters. As depicted in Figure 3D, TREVI^XMBD^’s Silhouette coefficients are notably higher than those of other methods. TREVI^XMBD^ excels in characterizing these temporal variations, maintaining within-cluster compactness, and enhancing between-cluster distinctions.

The above results corroborate that TF activities inferred by TREVI^XMBD^ effectively capture temporal variations among cells after perturbation and minimize the impact of non-biological signals.

### 3.4 TF activities improve the characterizations of cells from different batches

Batch effects represent a well-documented challenge in single-cell data analysis, exerting a detrimental impact on data integrity and precision [20-23]. Given the intrinsic nature of TF activities within individual cells, we hypothesized that incorporating TF activities could enhance the resilience of multi-omics data analysis against batch effects. To validate this hypothesis, we utilized the BMMC dataset, a comprehensive single-cell multi-omics dataset derived from 12 healthy human donors and produced as part of the NeurIPS 2021 Competition [10]. This dataset exhibited a complex array of nested batch effects stemming from variations among both donor identities and facility sites. In our experiment, we defined a ‘batch’ as cells obtained from a single donor and processed at a particular facility site, resulting in a total of 13 distinct batches.

As shown in Figure 4B, TREVI^XMBD^ obtained an average silhouette score of approximating 0.91 for the latent variables between batches (batch ASW), outperforming other methods. This observation suggests that TREVI^XMBD^ excels in the harmonization and amalgamation of cells originating from different batches. Figure 4C provides an overview of integration results for all methods, with each having three subplots indicating batch-based coloration, cell cluster-based coloration, and cell type-based coloration, respectively. Visually, TREVI^XMBD^’s approach displayed a homogeneous distribution of cells from diverse batches within the clusters, based on TF activities. In contrast, alternative methods exhibited the remaining batch-specific clusters. Take CD14 monocyte cells that are highlighted by red circles as an example, TREVI^XMBD^’s clustering effectively mitigated batch effects. Conversely, MultiVI separated CD14 monocyte cells into three clusters, each aligned with distinct batches. Similarly, Cobolt generated three clusters, while scMVP produced two clusters, suggesting the persistence of substantial batch effects in their results. Comparable outcomes were observed in the CD8 T cell population, marked by green circles.

**Figure 4.**
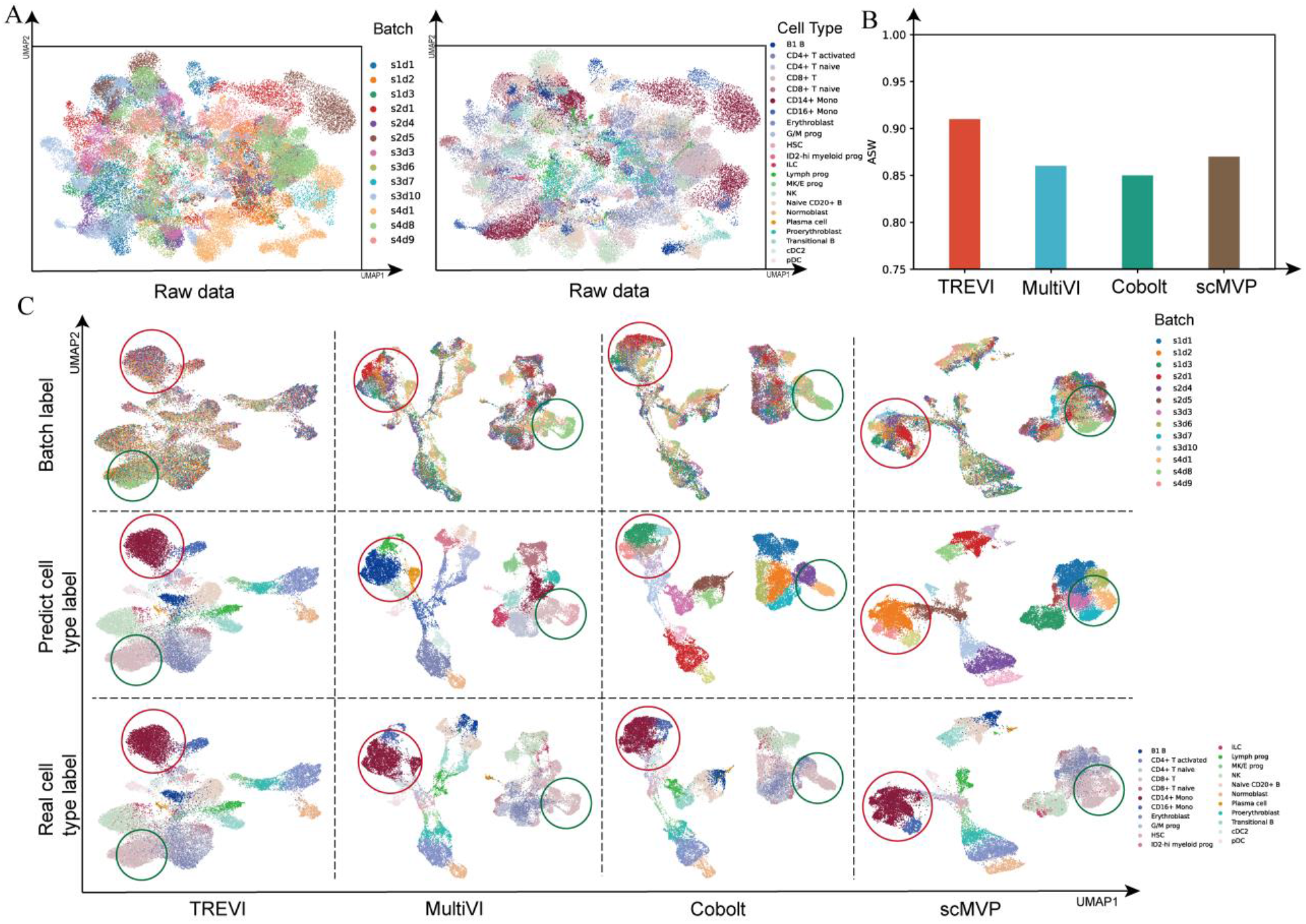
Performance comparison of batch effect correction. A. Before correction cells from the same cell type group by batches in UMAP clustering results of scRNA-seq data in the BMMC dataset, cells are colored by batch and cell type respectively. B. TREVI^XMBD^ and three other methods in batch ASW metric. C. After correction cells from the same cell type are mixed in clustering results on embedding of latent variables obtained by TREVI^XMBD^ and baseline methods in the BMMC dataset.

In summary, TREVI^XMBD^ effectively corrects batch effect, surpassing other multi-omics integration baseline models. TF activities robustly depict cells of the same types collected from various sources. TREVI^XMBD^ presents an effective solution to link data from different sources in a biologically meaningful manner to support the intended analysis [24].

### 3.5 TREVI^XMBD^’s generalization ability

TREVI^XMBD^ can serve as a pre-trained model suitable for application to new datasets, particularly those originating from the same tissue or featuring similar cell types, even when comprising scRNA-seq exclusively. To validate this, we trained TREVI^XMBD^ using the paired PBMC dataset and applied it to the PBMC-IFNB dataset. Initially, we conducted a visual inspection of the characteristics of the PBMC-IFNB dataset, revealing variations among cells within the same cell clusters or sharing identical cell types due to differences between two different batches (Figs. 5A-C). We then employed the pre-trained TREVI^XMBD^ to predict the cell-type identity for each cell in the PBMC-IFNB dataset and calculate low-dimensional cell embeddings, based on TF activities. The visualization of these cell embeddings demonstrated effective mitigation of batch-related variations.

**Figure 5.**
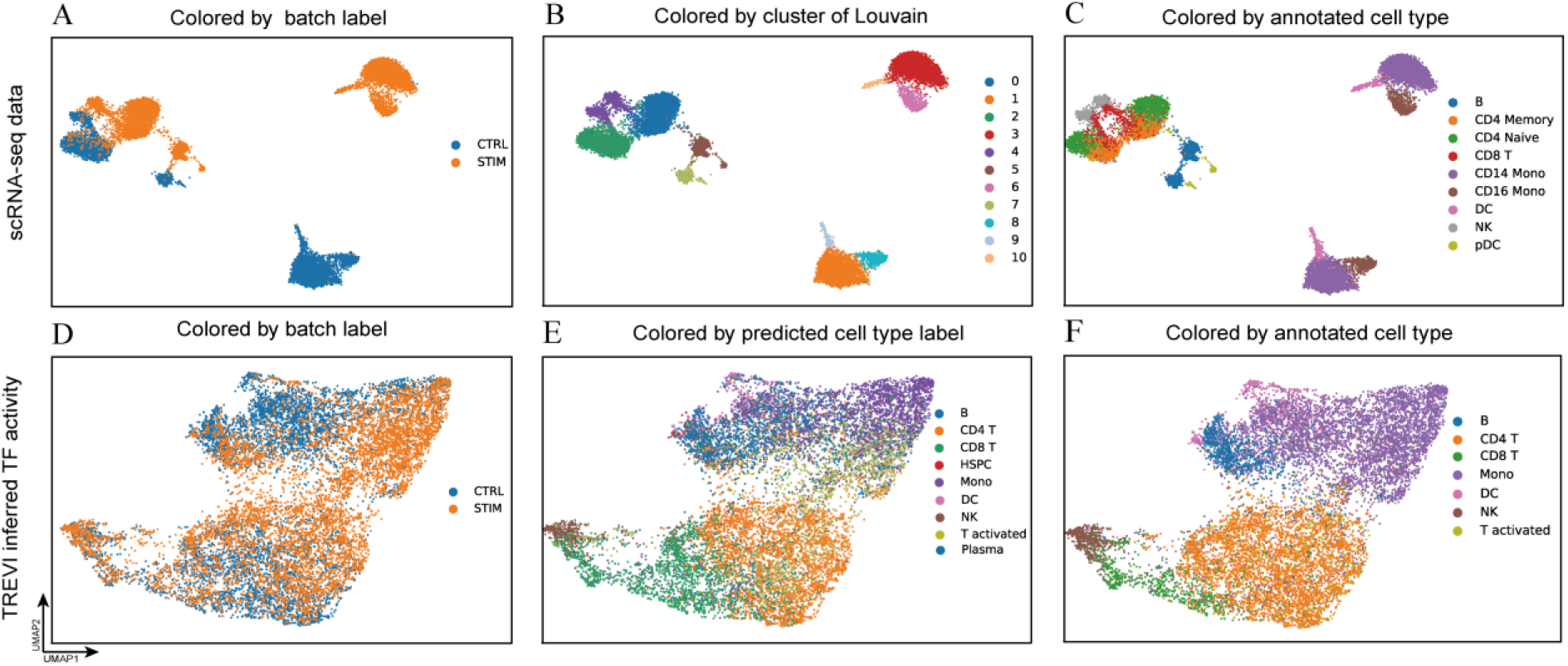
Performance of TREVI^XMBD^ as pre-trained model on the PBMC-IFNB dataset. The dataset is visualized in UAMP based on the scRNA-seq and the learned TF activity embeddings, and colored by batches, predicted cell types, and original cell type annotations, respectively.

Furthermore, the cell type predictions generated by TREVI^XMBD^ exhibited a high consistency with the cell type annotations provided by the data generator. Specifically, cells with the same predicted cell types aggregated closely (Fig. 5D-E) and occupied proximities congruent with the original cell type annotations (Fig. 5E-F). The results imply that after being pre-trained on datasets derived from multiple tissues, TREVI^XMBD^ has the potential to achieve atlas-level cell annotation.

### 3.6 TREVI^XMBD^ detects cell-specific TF-gene regulatory weights

TREVI^XMBD^ leverages established TF-gene regulatory relationships from publicly available databases as priors, and optimizes these regulations through Bayesian inference using the multi-omics data. Consequently, it is anticipated to unveil the dynamics of TF-gene regulation.

To assess this capability, we applied TREVI^XMBD^ to the PBMC dataset to investigate variations in TF-gene regulatory weights across distinct cell types. Figure 6A illustrates the inferred regulatory weights from the transcription factor E2F2 to nine of its target genes in three distinct cell types, namely CD14 monocyte cells, CD4 naïve T cells, and memory B cells. Notably, the regulatory weights of E2F2 exhibit variations across three cell types. Taking the target gene SPl1 as an example, E2F2 exhibits the highest regulatory weight in CD14 monocyte cells, a moderate weight in memory B cells, and the weakest weight in CD4 naïve T cells. Figure 6B displays the dynamics of E2F2’s regulatory weights to the nine target genes within individual cell types, emphasizing the pronounced changes in regulatory weights among cell types.

**Figure 6.**
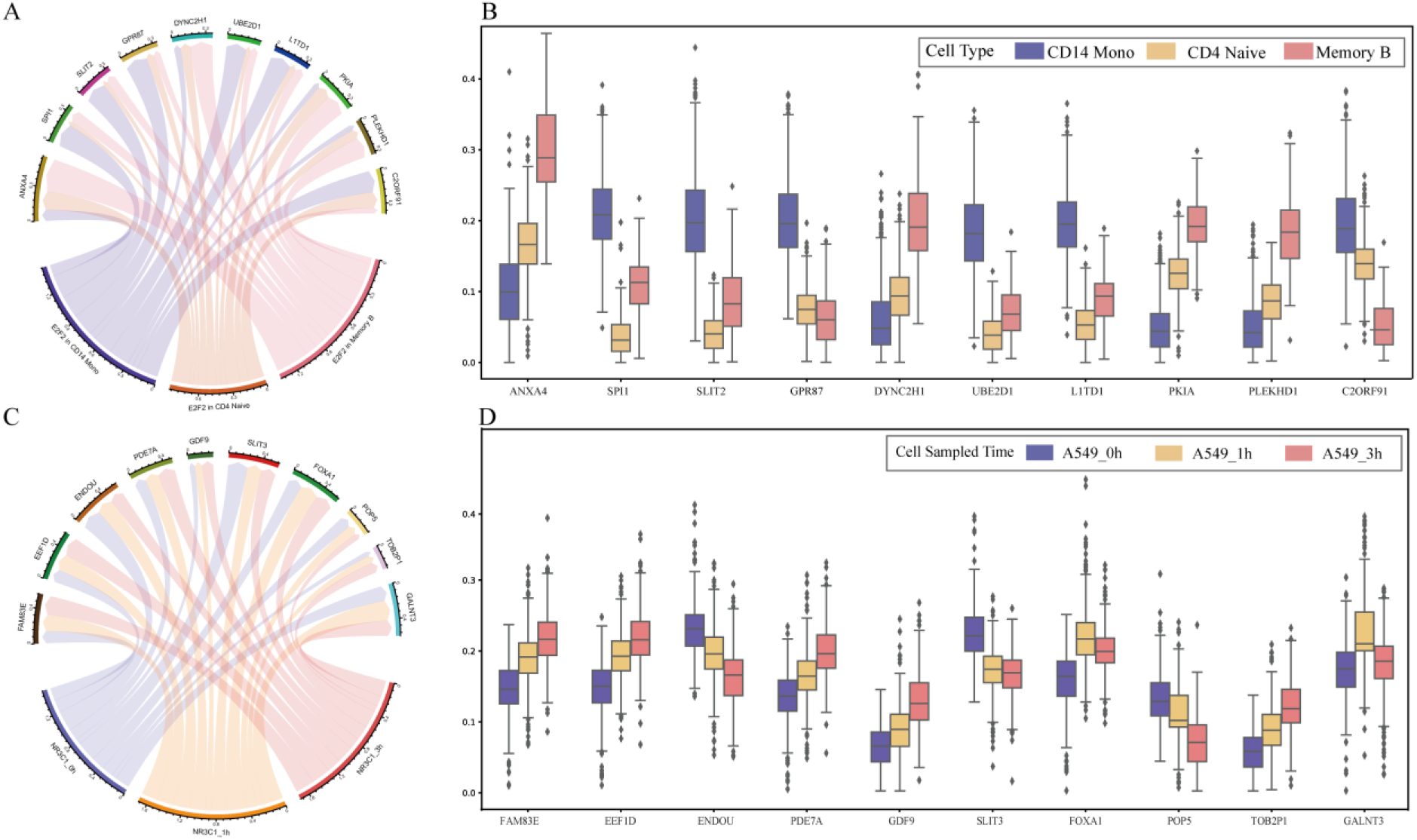
TREVI^XMBD^ infers specific TF-gene regulatory weights. A. Regulatory weights of E2F2 and its target genes across CD14 Mono, CD4 Naive, Memory B cells. B. Boxplot of regulatory weights between E2F2 and its target genes. C. Regulatory weights between NR3C1 and its target genes changes across times. D. Boxplot of regulatory weights between NR3C1 and its target genes.

Subsequently, we employed TREVI^XMBD^ to the A549 dataset to explore variations in the TF-gene regulatory weights throughout cell evolution. Figure 6C presents temporal fluctuations in the regulatory weights between the transcriptional factor NR3C1 and ten of its target genes. For instance, the regulatory weights between NR3C1 and its target gene ENDOU decreased over time after DEX treatment, while the regulatory weights between NR3C1 and its target gene FAM83E increased over time. Figure 6D displayed the dynamics of NR3C1’s regulatory weights to the ten target genes over time, highlighting substantial changes in regulatory weights over time were observed. In this experiment, TF-gene regulatory weights inferred by TREVI^XMBD^ not only infer cell type-specific gene regulatory relationships but also capture the dynamic changes of TF-gene regulation over time.

## 4. Discussion

The gene regulation study has witnessed significant advances with the advent of single-cell multi-omics sequencing. Single-cell multi-omics data cannot directly quantify cellular regulatory relationships, but it offers insights into molecular components during the regulatory processes, opening avenues to explore regulatory mechanism. Previous studies generally integrated multi-omics data to map them to a common embedding vector, overlooking the underlying biological mechanisms. Therefore, we propose TREVI^XMBD^, a Bayesian model that explicitly incorporates the generative process of gene expression profiles regulated by gene accessibility, TF activity and TF-gene regulatory weight. Our experiments demonstrate that TF activity inferred by TREVI^XMBD^ remains robust against batches, signal noise and technical bias, making it a valuable tool for characterizing cell types, tracking cell development, and removing batch effects. Moreover, TREVI^XMBD^ quantifies the regulatory strength of TFs on their target genes to construct a dynamic TF-gene regulatory network of single-cell resolution, providing valuable insights into the study of diverse regulatory patterns. After all, we aim to enhance the modeling capability of TREVI^XMBD^ by incorporating additional information, such as directed reference regulatory relationships or DNA methylation. Our primary objective is not only achieving high performance in specific tasks but also unveiling valuable insights within multi-omics data at the cellular level. We aspire to empower researchers with TREVI^XMBD^ to unravel the intricate regulatory mechanisms.

## Acknowledgements

Shaorong Fang and Tianfu Wu from Information and Network Center of Xiamen University are acknowledged for the help with high performance computing (HPC).

