## Supplementary material for "TREVI^XMBD^: A Transcriptional Regulation-driven Variational Inference Model to Speculate Gene Expression Mechanism with Integration of Single-cell Multi-omics": Supplymentary Materials (from another version of this preprint: version 2, 2023-11-25)

∗ To whom correspondence should be addressed:

### Supplementary materials S1. Data preprocessing

**Highly variable gene selection for scRNA-seq data:** The top 3000 highly variable genes (HVGs) are selected using *Scanpy* [1] (function *scanpy.pp.highly_variable_genes* with default parameters). Specifically, it normalizes the dispersions based on the gene expression values for each gene, and sorts them to select the highly variable genes. The single cell expression profile of top $G$ HVGs from $N$ cells are denoted as $\boldsymbol{X}$ and $\boldsymbol{X}_i$ is the gene expression profile of cell $i$.

**Chromatin accessibility scoring on scATAC-seq data:** Instead of using general cell-by-peak binary matrix, cell-by-gene chromatin accessibility scoring matrix is obtained using MAESTRO [2], a computational workflow for single-cell RNA-seq and ATAC-seq data. Specifically, MAESTRO designs a gene-by-cell matrix $\boldsymbol{R}$ to score the gene regulatory potential, which is defined by a matrix multiplication for each gene in each cell, which is defined as:

$$\boldsymbol{R} = \boldsymbol{W}^{\boldsymbol{T}}\boldsymbol{B}$$

where $\boldsymbol{B}$ is the binary peak-by-cell matrix obtained from single-cell ATAC-seq, and $\boldsymbol{W}$ is the peak-by-gene matrix, reflecting the distance between the center of peak and the transcription start site of gene. The regulatory score matrix $\boldsymbol{R}$ is correspondingly trimmed to the same dimension as $\boldsymbol{X}$, which are finally used as the paired inputs to TREVI$^{\text{XMBD}}$.

**TF-gene reference matrix building:** Following BITFAM developed by Gao et al. [3], the reference TF-gene regulating relationship i.e., the matrix $\boldsymbol{Ref}_{TF-gene}$, is obtained from the Gene Transcription Regulation Database (GTRD v19.04) [4], and $\boldsymbol{Ref}_{ij}$ uses 1 or 0 to indicates whether the database has recorded a regulatory relationship between TF $i$ and gene $j$.

### Supplementary materials S2. Loss function and parameter optimization

TREVI$^{\text{XMBD}}$ is a Bayesian-based probability model where all nonlinear transformations are achieved by neural networks and the parameters are optimized through variational inference with *Adam* Optimizer [5]. To approximate the true distribution $p_\theta(ls, y, rp, z_{TF}, z_{GRN}, z_y)$, the variational distribution is defined as $q_\varphi(ls, y, rp, z_{TF}, z_{GRN}, z_y|X)$, which can be decomposed as follows:

$$q_\varphi(ls, y, rp, z_{TF}, z_{GRN}, z_y|X) = q_\varphi(ls|X)q_\varphi(rp)q_\varphi(z_{TF}|X)q_\varphi(y|z_{TF})q_\varphi(z_y|y, z_{TF})q_\varphi(z_{GRN}|X)$$

Variational inference replaces the intractable marginal likelihood $p_\theta(x)$ with variational lower bound, and estimates the intractable integrals with low-variance Monte Carlo estimates, i.e., the reparameterization, which enables effective maximization of the evidence lower bound (ELBO). As a result, the data generated by the latent variables can be accessible to the real data by maximizing ELBO. In TREVI$^{\text{XMBD}}$, the transformation of maximizing the log-likelihood into maximizing the ELBO through variational inference can be formulated as follows:

$$\begin{aligned} log(p(X)) &= log\left(\int p_\theta(X|ls, y, rp, z_{TF}, z_{GRN}, z_y)p_\theta(ls, y, rp, z_{TF}, z_{GRN}, z_y)\, \boldsymbol{d}(ls, y, rp, z_{TF}, z_{GRN}, z_y)\right) \\ &= D_{KL}(q_\varphi(z_{TF}, z_{GRN}|x)||p_\theta(z_{TF}, z_{GRN}|X)) + L_{ELBO}(\theta, \varphi; X) \\ &\geq L_{ELBO}(\theta, \varphi; X) \end{aligned}$$

where

$$\begin{aligned} L_{ELBO}(\theta, \varphi; x) = \; & E_{q(z_{TF}, z_{GRN}|x)}\left[log\left(p_\theta(X|ls, y, rp, z_{TF}, z_{GRN}, z_y)\right)\right] \\ & - D_{KL}(q_\varphi(z_{TF}, z_{GRN}|X)||p_\theta(z_{TF}, z_{GRN})) \end{aligned}$$

$L_{ELBO}$ is composed of expected data likelihood, i.e., the reconstruction loss, and the KL divergence between the prior distribution and the constructed posterior distribution. To train the cell type classification and constrain the parameters when inferring $z_{GRN}$ within the same cell type, classification loss, denoted by $L_{Classify}$, which measures the distance between known cell types and predicted cell types using cross-entropy, is added to the overall loss function of TREVI$^{XMBD}$, as follows:

$$L = L_{ELBO} + \gamma L_{Classify}$$

$$= E_{q(z_{TF}, z_{GRN}|x)}\left[log\left(p_\theta\left(X|ls, y, rp, z_{TF}, z_{GRN}, z_y\right)\right)\right] - D_{KL}\left(q_\varphi(z_{TF}, z_{GRN}|x) \| p_\theta(z_{TF}, z_{GRN})\right)$$

$$+\gamma\left(-E\left[log\left(q_\varphi(y|z_{TF})\right)\right] - E\left[lo\,g\left(q_\varphi(y|z_{GRN})\right)\right]\right)$$

where γ is a hyperparameter that controls the weight of the classification loss in the overall loss.

### Supplementary materials S3. Table 1

**Table 1** The information of seven datasets for performance evaluation

| Datasets | No. of cells | No. of genes | No. of peaks | No. of cell types | No. of TF | Sequencing Technology | Source | Species |
|---|---|---|---|---|---|---|---|---|
| A549 | 3260 | 2494 | 189,603 | 3 | 34 | sci-CAR | Cao2018[6] | Human |
| Skin | 32,231 | 21,478 | 340,341 | 22 | 71 | SHARE-seq | Ma2020[7] | Mouse |
| Cerebral cortex | 9190 | 28,930 | 241,757 | 22 | 53 | SNARE-seq | Chen2019[8] | Mouse |
| PBMC | 9631 | 29,095 | 107,194 | 19 | 98 | 10x Multiome | Cao2022[9] | Human |
| BMMC | 69,249 | 129,921 | 116,490 | 22 | 97 | 10x Multiome | NIPS2021[10] | Human |
| Lymph node | 7039 | 3000 | 108,338 | 9 | 77 | 10x Multiome | Li2022[11] | Human |
| PBMC-IFNB | 12,687 | 14,053 | - | 9 | 98 | 10x Genomics | Kang2018[12] | Human |

### Supplementary materials S4. Evaluation metrics

When evaluating the performance of clustering and classification, we utilized the Adjusted Rand Index (ARI) and purity. The ARI is an external evaluation measure that quantifies the similarity between clustering results and actual labels. It improves upon the Rand Index (RI) by addressing the issue of RI's inability to ensure that randomly assigned clustering results have an RI value that close to zero. The ARI is calculated using the following formula:

$$\text{ARI} = \frac{(\text{RI} - \text{E[RI]})}{(\max(\text{RI}) - \text{E[RI]})}$$

Here, RI represents the Rand Index, E[RI] represents the expected Rand Index under random conditions, which is the average value we expect the Rand Index to have if the clustering results are random. The ARI ranges between -1 and 1, where a value closer to 1 indicates greater similarity in the clustering results, a value closer to -1 indicates dissimilarity in the clustering results, and a value close to 0 suggests clustering results that resemble random partitioning.

Purity is another commonly used evaluation metric for assessing the accuracy of clustering results. It computes the proportion of the most frequently occurring class within each cluster and sums these proportions. Purity ranges between 0 and 1, with a value closer to 1 indicating more accurate clustering results.

The purity is calculated using the formula:

$$\mathrm{Purity} = \frac{1}{N}\sum_{j=1}^{k} \max_{j} \left|w_{k} \cap c_{j}\right|$$

Here, N represents the total number of samples, $\Omega=\{w1, w2, \dots, wk\}$ denotes the cluster partition, and $C=\{c1, c2, \dots, cj\}$ represents the true class partition. The indices k and j refer to clusters and classes, respectively.

To evaluate the performance of removing batch effects, we employed the batch average silhouette width (ASW). ASW is an evaluation metric that measures the Euclidean distance and clustering results between samples from different batches. A higher ASW value indicates that the clustering results after removing batch effects exhibit better compactness and separability, meaning that cells of the same cell type from different batches are more tightly clustered. By utilizing the ASW metric, we can assess the performance of batch effect removal, determine whether the clustering quality of the processed data has improved, and ensure that batch effects do not excessively influence the results during further analysis and interpretation.

$$\mathrm{ASW} = \frac{1}{\mathrm{N}}\sum \frac{\left(\mathrm{b(i)} - \mathrm{a(i)}\right)}{\max\left(\mathrm{a(i)}, \mathrm{b(i)}\right)}$$

Here, N represents the number of samples, $\sum$ denotes the summation symbol, $b(i)$ represents the inter-batch distance for sample i, $a(i)$ represents the intra-batch distance for sample i, and $\max(a(i), b(i))$ represents the larger of $a(i)$ and $b(i)$.

### Supplementary materials S5. Baseline methods

MultiVI [13] is a multi-modal variational inference method that uses variational autoencoders to analyze and integrate single-cell multi-omics data through probabilistic modeling, aims to learn latent variables containing shared information between the two modes from the multi-omics data and realize cross-modal imputation.

scMVP [14] (single-cell Multi-View Profiler) is a multi-modal deep generative model designed for handling sequencing data that simultaneously measure gene expression and chromatin accessibility in the same cell, scMVP combines transformer and multimodal variational autoencoder, generates common latent representations for dimensionality reduction and cell clustering.

Cobolt [15] employs a hierarchical Bayesian generative model-based multimodal variational autoencoder to extract shared latent variables that capture shared information from multi-omics data, utilizing joint-modality data to enable the joint analysis of cells sequenced on separate sequencing modalities.
